# Force-profile analysis of the cotranslational folding of HemK and filamin domains: Comparison of biochemical and biophysical folding assays

**DOI:** 10.1101/470831

**Authors:** Grant Kemp, Renuka Kudva, Andrés de la Rosa, Gunnar von Heijne

## Abstract

We have characterized the cotranslational folding of two small protein domains of different folds – the α-helical N-terminal domain of HemK and the β-rich FLN5 filamin domain – by measuring the force that the folding protein exerts on the nascent chain when located in different parts of the ribosome exit tunnel (Force-Profile Analysis - FPA), allowing us to compare FPA to three other techniques currently used to study cotranslational folding: real-time FRET, PET, and NMR. We find that FPA identifies the same cotranslational folding transitions as do the other methods, and that these techniques therefore reflect the same basic process of cotranslational folding in similar ways.

---

In recent years, a number of new experimental methods for analyzing the cotranslational folding of protein domains have been developed. These include real-time FRET analysis [1], and methods in which nascent polypeptide chains of defined lengths are arrested in the ribosome and their folding status analyzed by, e.g., cryo-EM [2, 3], protease resistance [1, 4-9], NMR [10-15], photoinduced electron transfer (PET) [1, 16], folding-associated co-translational sequencing [17], optical tweezer pulling [18-21], fluorescence measurements [22], and measuring the force that the folding protein exerts on the nascent chain using a translational arrest peptide (AP) as a force sensor [2, 3, 9, 21, 23-25]. Further, coarse-grained molecular dynamics simulations of various flavors can provide detailed insights into cotranslational folding reactions [26-30], especially when coupled with experimental studies [3, 26, 31].

While these different methods have all been used with success, they have rarely been directly compared on the same target protein. As a first step towards such comparisons, we now report AP-based Force-Profile Analysis (FPA) for two protein domains - the α-helical HemK N-terminal domain (HemK-NTD) and the β-sheet-rich FLN5 filamin domain - that have previously been studied by real-time FRET/PET, and NMR respectively [1, 14, 16]. We find that FPA identifies the same cotranslational folding transitions as do the other methods. We conclude that results obtained with the different methods can be interpreted under a common conceptual framework, albeit at different levels of detail and with different limitations inherent to the analysis.

## AP-based force-profiles

FPA is based on the observation that the efficiency of AP-induced translational stalling is reduced when the nascent chain is subject to an external pulling force [21, 32, 33]. As shown in Fig. 1, such a pulling force can be induced by cotranslational protein folding. For constructs where the tether+AP segment is long enough that there is just enough space in the ribosome exit tunnel for the protein to fold if the segment is stretched out from its equilibrium length, some of the free energy gained upon folding will be stored as elastic energy (increased tension) in the nascent chain, reducing stalling and increasing the relative amount of full-length protein (Fig. 1a). By measuring the stalling efficiency (measured as fraction full-length protein, *f*_*FL*_) for a series of constructs of increasing tether length (Fig. 1b, c), a force profile can be generated that shows how the folding force varies with the location of the protein in the exit tunnel [2], and hence when during translation the protein starts to fold. FPA has been used to map the cotranslational folding of more than 10 different proteins so far [9].

**Figure 1.**
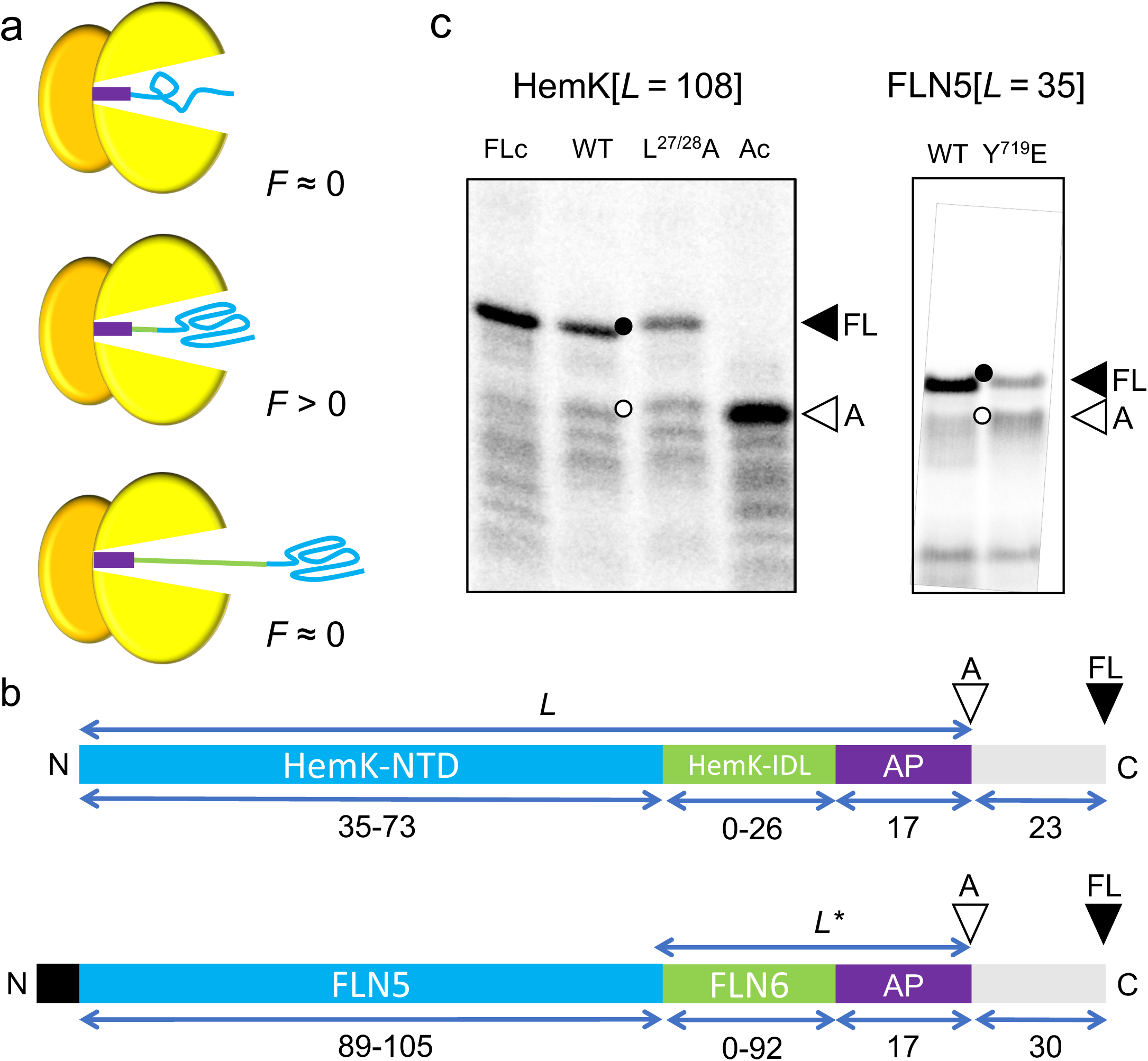
The Force Profile Analysis (FPA) assay. (a) Cotranslational folding can generate force on the nascent chain. The nascent protein is in blue, the tether in green, the AP in purple, and the ribosomal subunits in yellow. At short tether lengths, the protein is still too deep in the exit tunnel to fold when the AP is reached, there is negligible force exerted on the stalled nascent chain (*F* ≈ 0, top panel), and the arrested (*A*) form of the protein is the main product. Similarly, at long tether lengths the protein is already folded when the AP is reached, little force is exerted (*F* ≈ 0, bottom panel), and mostly *A* is produced. However, at some intermediate tether length when there is just enough space in the exit tunnel for the protein to fold if the tether is stretched, force will be exerted on the nascent chain (*F* > 0, middle panel), reducing stalling and increasing the amount of full-length (*FL*) product. (b) Schematic representation of the HemK and FLN5 constructs (full sequences can be found in Supplementary Table 1). HemK-NTD (and its truncations) or FLN5 (blue) were tethered to the SecM(*Ec*) AP (purple) by a varying-length part of their respective native downstream sequence (green), either the HemK interdomain linker (HemK-IDL) or the FLN6 filamin domain. Following the AP, a sequence of 23 (for HemK) or 30 (for FLN5) amino acids derived from LepB is included to allow the resolution of arrested (*A*) or full length (*FL*) products by SDS-PAGE (empty and filled triangles, respectively). The HemK-IDL was progressively truncated from its C-terminal end from 26 down to zero residues, followed by truncation of HemK-NTD from 73 down to 35 residues, as indicated by the numbers underneath the cartoon representations. Likewise, the FLN6 linker segment was truncated from its C-terminal end from 92 down to zero residues, followed by truncation of FLN5 from 105 down to 89 residues. (c) SDS-PAGE of the *in vitro* translation products of the HemK[*L* = 108] (left) and FLN5[*L*=35] (right) with arrested (*A*) and full-length (*FL*) products indicated. HemK[*L* = 108]: lane 1, full length control (FLc) construct where the C-terminal, critical Pro of the SecM(*Ec*) AP is substituted by Ala, which eliminates arrest; lane 2, wild-type construct; lane 3, unstable mutant L^27^A+ L^28^A; lane 4, arrest-control (Ac) construct where the codon immediately following the AP has been mutated to a TAA stop codon. FLN5[*L*=35]: lane 1, wild-type FLN5[*L*=35] construct; lane 2, non-folding mutant control Y^719^E. Materials and Methods are available as a Supplement.

## Cotranslational folding of the HemK N-terminal domain

In a pioneering study, Holtkamp *et al.* followed the cotranslational folding of the 73-residue long HemK-NTD (Fig. 2a) in real time using both FRET- and PET-based assays [1]. A subsequent, more detailed analysis [16] led to the identification of four folding intermediates (I-IV) in addition to the native state (V), which were proposed to correspond to the linear, stepwise addition of individual α-helices to a growing, compact core. Furthermore, it was concluded that the folding reaction is rate-limited by translation (which proceeds at a rate of ∼4 codons s^−1^ in the *in vitro* translation system used [1]). A coarse-grained molecular dynamics study has also provided evidence for the cotranslational appearance of compact folding intermediates composed of the first three (intermediate II) and first four (intermediate III) α-helices of HemK-NTD, followed by de-compaction (intermediate IV) and folding into the native 5-helix bundle (V) at longer tether lengths [26].

**Figure 2.**
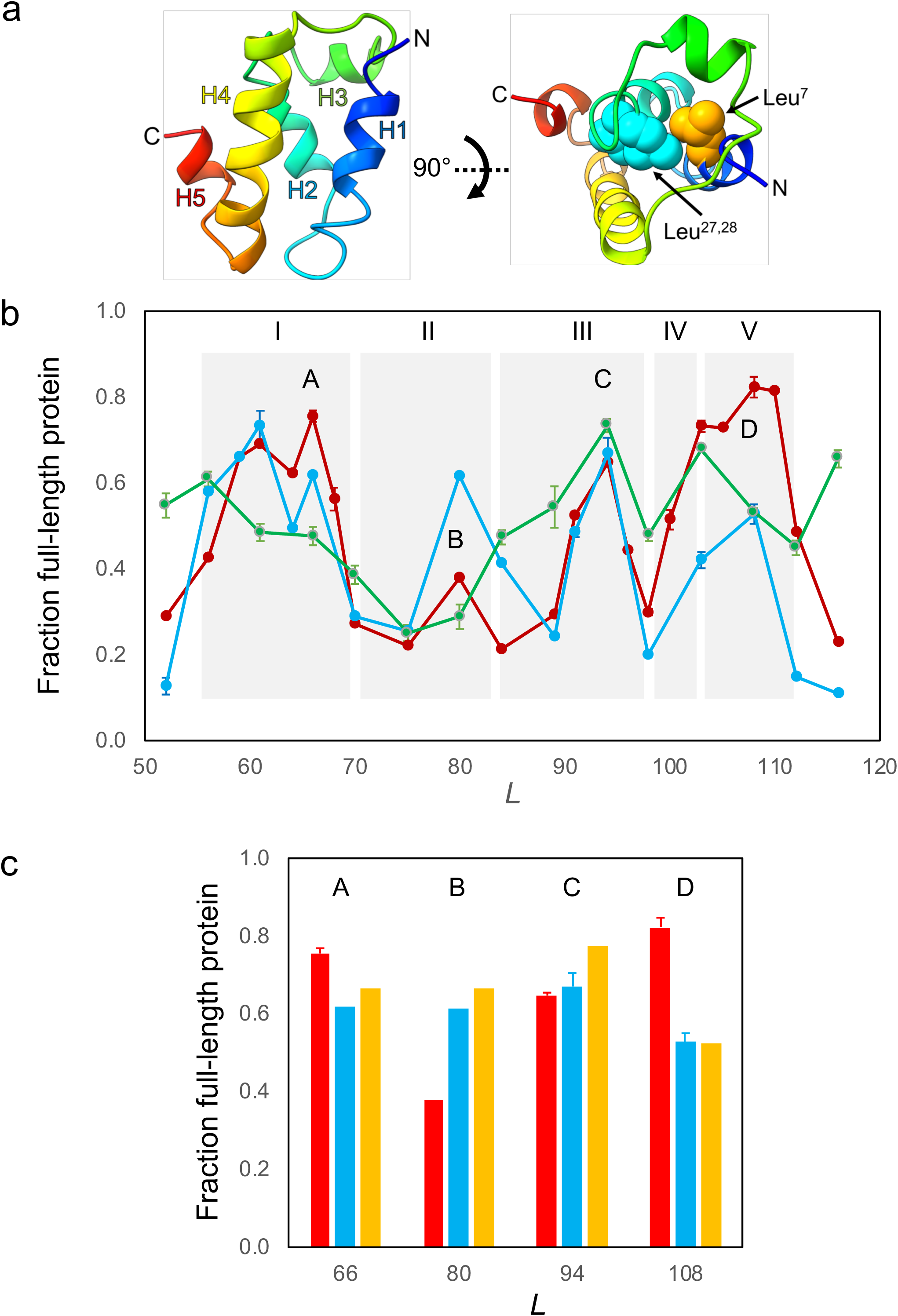
FPA of Hemk-NTD using the SecM(*Ec*) AP as the force sensor. (a) Ribbon representation of the backbone structure of HemK-NTD (PDB 1T43 [40]) rainbow-colored from the N-terminus (blue) to the C-terminus (red) showing the five a-helices (H1-H5). Leu^7^(orange), Leu^27^, and Leu^28^ (blue) are shown as spheres. (b) Force profiles obtained by translation in the *E. coli*-derived PURE *in vitro* transcription-translation system (red curve), by *in vivo* expression in *E. coli* (green curve), and by *in vitro* expression of the unstable L^27,28^A mutant (blue curve). Peaks in the force profile discussed in the text are labelled A-D. The shaded areas labelled I-V indicate the approximate *L*-values corresponding to the folding intermediates (I-IV) and the native structure (V) derived from real-time FRET/PET measurements [1, 16]. *L* is the number of residues between the N-terminal end HemK-NTD and the C-terminal residue of the AP (c.f., Fig. 1b). (c) *f*_*FL*_ values for wild-type HemK-NTD (red), the L^27^A+ L^28^A mutant (blue), and the L^7^N (orange) mutant at *L* = 66, 80, 94, and 110 residues (corresponding to peaks A-D in the force profile in panel *a*, as indicated). Error bars indicate SEM values.

In order to compare FPA to the FRET and PET assays, we recorded a force profile for HemK-NTD at ∼5-residue resolution using the *Escherichia coli* SecM(*Ec*) AP, Fig. 2b (red curve; in this case, in order to be consistent with the previous publications [1, 16], we measured *L* from the N-terminus of HemK-NTD to the C-terminal end of the AP, Fig. 1b). Four peaks (A-D) are apparent in the force profile. Peak D (*L* ≈ 100-110 residues) coincides with the formation of the native state V as seen by FRET, and corresponds to a situation where the C-terminal end of HemK-NTD is 30-35 residues away from the ribosome peptide transferase center (PTC), close to the exit-port region where other proteins of similar size have been found to fold into their native state [3, 24]. Strikingly, peaks A-C in the force profile nicely match the folding intermediates I-III identified by real-time FRET and PET analysis [16], although peak B is just barely detectable above the background. The proposed intermediate IV is not resolved in the force profile, but may contribute to the early shoulder of peak D.

To test how mutations in the hydrophobic core of HemK-NTD affect the observed forces, we recorded a force profile for a mutant HemK-NTD in which Leu^27^and Leu^28^ in helix H2 were simultaneously mutated to Ala (Fig. 2a, and blue curve in Fig. 2b). The L^28^A mutation reduces the *in vitro* denaturation temperature of HemK-NTD from 50°C to 30°C [1]. The [L^27^A+ L^28^A] mutation caused a strong reduction of the amplitude of peak D, but had only marginal effects on peaks A and C. Interestingly, the mutation led to a marked increase the amplitude of peak B. To confirm these effects, we also measured *f*_*FL*_ values for a Leu^7^ to Asn mutation in helix H1 (Fig. 2a) for *L* values corresponding to peaks A-D. As seen in Fig. 2c, the L^7^N mutation behaves similarly to the [L^27^A+ L^28^A] mutation: *f*_*FL*_ is strongly reduced for peak D, not affected for peaks A and C, and increased for peak B.

Finally, we also recorded an *in vivo* force profile by expression of the HemK-NTD constructs in *E. coli* cells. An HA tag was added just before the AP to enable the detection of HemK-NTD translation products by immunoprecipitation. Guided by previous mutagenesis studies of a SecM AP [34], we placed the HA tag overlapping the SecM(*Ec*) AP, resulting in an HA-AP sequence that was of the same length and somewhat lower stalling strength than the SecM(*Ec*) AP used in the *in vitro* experiments (Supplementary Fig. S1). The HemK-NTD force profile obtained *in vivo*, Fig. 2b (green curve) is similar to the one obtained *in vitro* (Fig. 2b, red curve), although peaks A and D are shifted to somewhat lower *L* values. Considering the lack of chaperones such as trigger factor and GroEL/ES in the PURE *in vitro* translation system and the ∼20-fold faster translation rate *in vivo*, the correspondence between the *in vitro* and *in vivo* force profiles is remarkably good.

We conclude that FRET/PET, MD, and FPA all give similar a similar picture of the cotranslational folding of HemK-NTD, and all detect the formation of at least two folding intermediates as well as the native state. Mutation of residues in the hydrophobic core of HemK-NTD reduce the amplitude of peak D (corresponding to the native state) in the force profile as expected, but have little effect on peaks A and C and even increase the amplitude of peak B, suggesting that the corresponding folding intermediates may have less well-defined tertiary structures than the native fold and hence be less sensitive to individual point-mutations.

## Cotranslational folding of the FLN5 domain

The cotranslational folding of the Ig-like FLN5 filamin domain has previously been studied by NMR measurements on ribosome-attached nascent chains and of purified C-terminally truncated versions of the protein [13, 14]. The main conclusion from these studies is that FLN5 folds only when it has fully cleared the exit tunnel and is some distance away from the ribosome surface, at a tether length of 40-45 residues. This tether length is clearly longer than what is required for folding of a similar-sized Ig-like domain such as the I27 titin domain that folds at a tether length of ∼35 residues, as determined by FPA and cryo-EM [3]. The simplest explanation for the long tether length required for cotranslational folding of FLN5 is that the ribosome surface destabilizes the folded state relative to the unfolded state [14], as has been observed for other proteins when tethered to a ribosome [20, 35, 36].

Given that the NMR data indicate that FLN5 folds at a considerably longer tether length than I27, we were curious to see if this is also reflected in its force profile. The FLN5 force profile obtained with a mutant AP, SecM(*Ec*, S→K) (see Supplementary Materials and Methods), that is somewhat more resistant to pulling force than is SecM(*Ec*) is shown in Fig. 3, together with a curve derived from the NMR data showing averaged relative intensities of three ^15^N amide resonances arising from the unfolded FLN5 domain [14]. To be consistent with the published NMR data [13, 14], in this case we plot the data as a function of *L*^*\**^, i.e., the distance between the C-terminal end of the FLN5 domain and the PTC (c.f., Fig. 1b). The force profile has two peaks: one at *L*^*\**^ = 35-47 residues that coincides with the main folding transition detected by NMR, and one at *L*^*\**^ ≈ 20 residues. There is an apparently significant dip both in the force profile and the NMR profile at *L*^*\**^ = 43 residues; at present, we have no explanation for this precisely localized feature in the two profiles.

**Figure 3.**
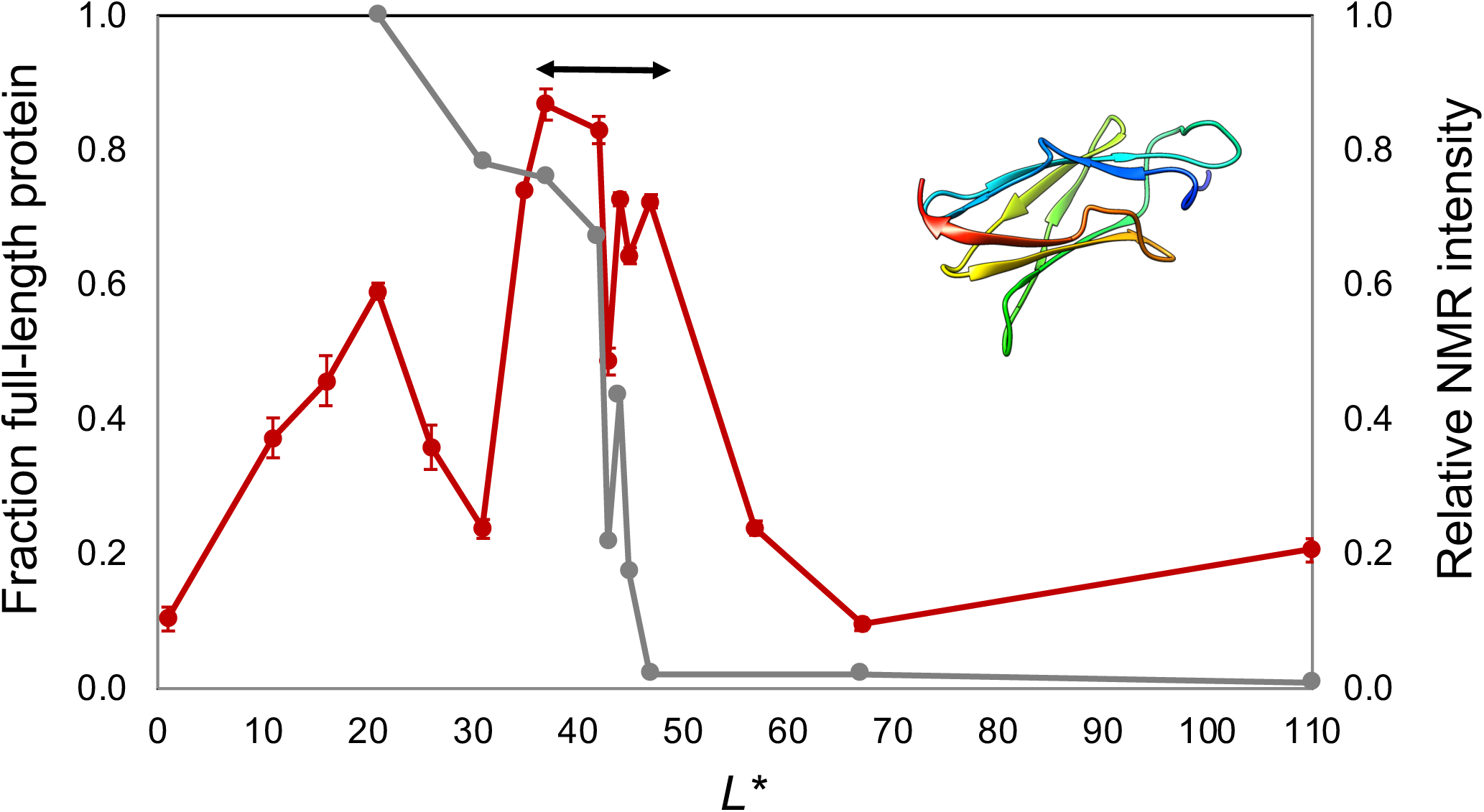
FPA of FLN5 using the SecM(*Ec*, S→K) AP as the force sensor (red curve) and averaged relative NMR intensities of three ^15^N amide resonances arising from the unfolded FLN5 domain (grey curve) [14] as a function of *L*^*\**^. The force profile was obtained by *in vitro* translation in the *E. coli*-derived PURE *in vitro* transcription-translation system. Error bars indicate SEM values. The arrow indicates the location of the folding transition derived from the NMR data. *L*^*\**^ is the number of residues between the C-terminal end of the FLN5 domain and the C-terminal residue of the AP (c.f., Fig. 1b). Inset: ribbon representation of the backbone structure of FLN5 (PDB 2N62), colored from N-terminus (blue) to C-terminus (red).

To validate that the main peak at *L*^*\**^ = 35-47 residues in the force profile reflects folding into the native state, we also analyzed an unstable mutant of FLN5, Y^719^E, using the weaker SecM(*Ec*) AP. As expected, the main peak is of significantly lower amplitude in this mutant; notably, however, the peak at *L*^*\**^ ≈ 20 residues is unaffected (Supplementary Fig. S2 blue curve), mirroring the insensitivity of the early peaks in the HemK-NTD force profile to the [L^27^A+ L^28^A] mutation (Fig. 2b).

The peak in the force profile at *L*^*\**^ = 35-47 residues is somewhat wider than the folding transition detected by NMR. Presumably, this is because the NMR resonances reflect only the folding transition itself, whereas the force profile reflects forces generated not only by the folding transition, but also by, e.g., the folded protein when tethered close to the ribosome surface [37]. In line with this, the peak in the force profile extends to higher *L*^*\**^ values when the weaker SecM(*Ec*) AP is used, Supplementary Fig. S2.

In conclusion, the main cotranslational folding transition of FLN5 to the native state is detected both by NMR and FPA, and a putative folding intermediate is also apparent in the FPA data.

## Summary

We have characterized the cotranslational folding of two small protein domains of different folds – the α-helical HemK-NTD and the β-rich FLN5 – by FPA, allowing us to compare FPA to three other techniques currently used to study cotranslational folding: real-time FRET, PET, and NMR. The results show broad agreement between FPA on the one hand, and the three biophysical techniques on the other. Previously, we have shown that FPA also agrees with results from on-ribosome pulse-proteolysis [9] and coarse-grained MD simulations [2, 3, 38].

It thus appears that all these techniques reflect the same basic process of cotranslational folding in similar, yet distinct ways. FPA differs from the other presently used techniques in that it reflects not only the cotranslational folding process *per se*, but in principle also other forces such as that expected to be generated when a folded protein is tethered close to the ribosome surface [37]. Our results nevertheless indicate that the methods discussed here can be expected to yield comparable results, at least for proteins that fold fast relative to the rate of translation. For proteins that fold more slowly, methods such as NMR, PET, pulse-proteolysis and FPA that are based on an analysis of nascent chains stably tethered to the ribosome may indicate folding at somewhat shorter tether lengths than will methods such as real-time FRET, in which the nascent chain is being continuously lengthened during the assay [39].

Techniques that use stable tethering can potentially yield folding profiles with single-residue resolution, since the nascent chain can be lengthened one residue at a time. Real-time techniques, in contrast, average over a population of elongating ribosome-nascent chain complexes and further depend on an estimate of the local translation rate to convert from time to nascent-chain length, and therefore cannot achieve single-residue resolution. It is still an open question as to what extent the fine structure of single-residue resolution data can yield useful information on the folding process.

It is particularly interesting to note that all the techniques appear able to detect intermediates on the cotranslational folding pathway, not just the formation of the native state. In-depth analysis of cotranslational folding intermediates may shed further light on how the presence of the ribosome affects protein folding.

## Acknowledgements

FLN5 constructs previously used for NMR studies were kindly provided by Lisa Cabrita and John Christodoulou, University College London. This work was supported by grants from the Knut and Alice Wallenberg Foundation, the Swedish Cancer Foundation, and the Swedish Research Council to GvH.

## Supplement 1

### Materials and Methods

#### Plasmids

All oligonucleotides for DNA manipulation were purchased from Eurofins Genomics. All enzymes were from New England Biolabs (NEB). All sequences were confirmed by DNA sequencing (Eurofins Genomics).

*HemK.* For translation in the PURExpress^®^ system (NEB), the entire HemK gene was amplified from *Escherichia coli* BL21(DE3) by colony PCR. The amino-terminal 99 amino acids of HemK were fused to the SecM arrest peptide from *Escherichia coli* (SecM(*Ec*) [1] followed by 69 nucleotides (corresponding to 23 amino acids) from the LepB, and the resulting gene was cloned gene a pET19b vector by Gibson Assembly [2]. Additional truncations were made by overlap extension PCR, shortening HemK in 2 to 4 amino acids steps until only 35 N-terminal amino acids of HemK remained. Substitutions in the sequence (Leu^7^Asn and Leu^27,28^Ala) were introduced by site-directed mutagenesis PCR. To allow immunodetection following *in vivo* translation, a hemagglutinin (HA) tag was substituted into the SecM sequence by overlap extension PCR yielding: 5’- <u>GGTAGCTACCCATACGATGTTCCAGATTACGCT</u>CCCATCCGTGCTGGCCCT-3’ (substituted sequence vs SecM is underlined) and the protein sequence <u>GS</u>**<u>YPYDVPDYA</u>**PIRAGP (HA tag in bold). The sequences with the HA-modified arrest peptide were subcloned into a pING vector [1] using Gibson assembly [2]. Sequences for all constructs are included in Supplementary Table S1.

*FLN5.* The previously described FLN5 constructs (21-110) fused to the SecM(*Ec*) arrest peptide [3] were engineered to include 93 nucleotides (corresponding to 30 amino acids) from the LepB gene at the 3’ end of the SecM(*Ec*) sequence, to enable identification of arrested and full-length variants in the force-profile assays. Additional C-terminal truncations in FLN5 (1-16) (see Supplementary Table 1) were generated by inverse PCR using phosphorylated primers and ligation.

The SecM(*Ec*, S→K) arrest peptide variant with increased stalling strength was generated by substituting the codon for serine at position -10 (relative to the critical C-terminal proline residue at the end of the AP) with the AAG codon for lysine, yielding the sequence 5′- TTCAGCACGCCCGTCTGGATA<u>AAG</u>CAGGCGCAAGGCATCCGTGCTGGCCCT- 3′, translated as the protein sequence FSTPVWI<u>K</u>QAQGIRAGP. Sequences for all constructs are included in Supplementary Table S1.

#### In vitro transcription and translation

HemK constructs were translated using the PURExpress^®^ kit (NEB) for 15 min at 37°C in the presence of [^35^S]-methionine (Perkin Elmer) from a linear portion of the plasmid DNA generated by PCR using T7 DNA sequencing primers. The translation reaction was stopped by adding a final concentration of 5% TCA to the mix and incubation on ice for 30 min. The TCA-precipitated samples were then centrifuged at 20,000x*g* for 10 min, solubilized in Laemmli buffer supplemented with 400 µg/ml RNAse A (Sigma), incubated at 37°C for 30 min, and subsequently analyzed by SDS-PAGE (16% Tricine gels from Thermo Fisher Scientific) and autoradiography. All *f*_*FL*_ measurements were carried out at least in duplicate (all pairwise differences in *f*_*FL*_ ≤ 0.06) and points where error bars are given (SEM) were repeated 3 or 4 times.

FLN5 constructs were translated in the PURExpress® kit for 20 min at 37°C from plasmid DNA (300 ng) in the presence of [^35^S]-methionine. The reaction was treated as above for HemK with the exception the 12% Bis-Tris gels from Thermo Fisher Scientific were used in place of 16% Tricine gels.

#### In vivo translation

HemK constructs in the pING vector were transformed into chemically competent MC1061 *E. coli* and selected on LB agar plates containing 100 µg/mL of carbenicillin. A single colony was selected and grown overnight in M9 minimal medium supplemented with 19 standard amino acids (1 mg/ mL) excluding methionine, 100 mg/mL thiamine, 0.1 mM CaCl_2_, 2 mM MgSO_4_, 0.4% (w/v) fructose, 100 µg/mL ampicillin at 37°C. The following day, the overnight culture was back-diluted to OD_600_ of 0.1 into fresh M9 medium and allowed to grow at 37°C until reaching an OD_600_ of 0.35. Expression was started by the addition of 0.2% arabinose. After 5 min, HemK was pulse-labelled for 2 min with [^35^S]-methionine. Expression was halted by the addition of 17% (w/v final concentration) of cold TCA and incubation for 30 min on ice. Following centrifugation (20,000x*g*, 4°C, 10 min) the supernatant was removed by aspiration and residual TCA and lipids were removed by addition of 0.5 mL of cold acetone followed by a repeat centrifugation and aspiration. The pellets were air-dried for a few minutes on the bench,then resuspended in 100 uL Tris-SDS buffer (10 mM Tris HCl pH 7.5, 2% SDS). The sample was precleared using 10 uL of protein G sepharose beads (GE Healthcare), washed with TSET buffer (20 mM Tris-HCl pH7.5, 150 mM NaCl, 1 mM EDTA, 0.1% Tween-20), and HemK was immunoprecipitated on 10 uL protein G sepharose beads pre-complexed with 1 uL of HA-antibody (Mouse IgG1, κ from Biolegend) for 1 hr at 4°C. Following two wash steps ([20 mM Tris-HCl pH7.5, 150 mM NaCl, 2 mM EDTA, 0.2% Tween-20] and [10 mM Tris-HCl pH 7.5]), HemK was eluted from the beads by the addition of 10 uL of Laemmli buffer supplemented with 400 µg/mL of RNAse I and processed in the same way as the *in vitro* translated HemK samples above. All points were measured in triplicate beginning with different colonies. Error bars indicate SEM values. SDS-PAGE gels of all constructs are included in Supplementary Figures S3-S6.

#### Quantification of radioactively labelled proteins

Intensity profiles of the protein bands on the gels were generated using MultiGauge (Fujifilm) or NIH ImageJ [4] and were fit to a Gaussian distribution using the in-house software EasyQuant (Rickard Hedman, Stockholm University). The area under the curves representing the arrested and full-length bands (*I*_*A*_, *I*_*FL*_) was determined and was used to estimate the fraction full-length protein for each construct, *f*_*FL*_ = *I*_*FL*_/(*I*_*FL*_ + *I*_*A*_). All quantifications are included in Supplementary Data 1.

### Molecular graphics

Images of protein structures were prepared in Chimera [5].

---

**Supplementary Table 1.** *In vitro* expression constructs of HemK: [HemK-amino terminal domain]-[HemK linker domain]-[Arrest Peptide]-[LepB derived C-terminal tail]. For *in vivo* expression, an HA tag was added to the Arrest Peptide to give the sequence GSYPYDVPDYAPIRAGP. Sequences where the Leu^27,28^Ala substitutions were made are <u>underlined</u> and those where the Leu^7^Asn was made are highlighted.

| Total distance to PTC (aa) | Translated protein sequence |
| --- | --- |
| 116 | MEYQHWLREAI SQLQASESPRRDAE <u>IL</u>LEHVTGKGRTFILAFGETQLTDEQC<br>QQLDALLTRRRDGEPIAHLTG VREFWSLPLFVSPATLIPRPDTECLVFSTPV<br>WISQAQGI RAGP GSSDKQEGEWPTGLRLSRIGGIH* |
| 112 | MEYQHWLREAI SQLQASESPRRDAE <u>IL</u>LEHVTGKGRTFILAFGETQLTDEQC<br>QQLDALLTRRRDGEPIAHLTG VREFWSLPLFVSPATLIPRPDTFSTPVWISQ<br>AQGI RAGP GSSDKQEGEWPTGLRLSRIGGIH* |
| 110 | MEYQHWLREAI SQLQASESPRRDAE <u>IL</u>LEHVTGKGRTFILAFGETQLTDEQC<br>QQLDALLTRRRDGEPIAHLTG VREFWSLPLFVSPATLIPRPFSTPVWISQAQ<br>GIRAGP GSSDKQEGEWPTGLRLSRIGGIH* |
| 108 | MEYQHW **L**REAI SQLQASESPRRDAE <u>IL</u>LEHVTGKGRTFILAFGETQLTDEQC<br>QQLDALLTRRRDGEPIAHLTG VREFWSLPLFVSPATLIPFSTPVWISQAQGI<br>RAGP GSSDKQEGEWPTGLRLSRIGGIH* |
| 105 | MEYQHWLREAI SQLQASESPRRDAE <u>IL</u>LEHVTGKGRTFILAFGETQLTDEQC<br>QQLDALLTRRRDGEPIAHLTG VREFWSLPLFVSPATFSTPVWISQAQGI RAG<br>P GSSDKQEGEWPTGLRLSRIGGIH* |
| 103 | MEYQHWLREAI SQLQASESPRRDAE <u>IL</u>LEHVTGKGRTFILAFGETQLTDEQC<br>QQLDALLTRRRDGEPIAHLTG VREFWSLPLFVSPFSTPVWISQAQGI RAGP G<br>SSDKQEGEWPTGLRLSRIGGIH* |
| 100 | MEYQHWLREAI SQLQASESPRRDAE <u>IL</u>LEHVTGKGRTFILAFGETQLTDEQC<br>QQLDALLTRRRDGEPIAHLTG VREFWSLPLF FSTPVWISQAQGI RAGP GSSD<br>KQEGEWPTGLRLSRIGGIH* |
| 98 | MEYQHWLREAI SQLQASESPRRDAE <u>IL</u>LEHVTGKGRTFILAFGETQLTDEQC<br>QQLDALLTRRRDGEPIAHLTG VREFWSL P FSTPVWISQAQGI RAGP GSSDKQ<br>EGEWPTGLRLSRIGGIH* |
| 96 | MEYQHWLREAI SQLQASESPRRDAE <u>IL</u>LEHVTGKGRTFILAFGETQLTDEQC<br>QQLDALLTRRRDGEPIAHLTG VREFWS FSTPVWISQAQGI RAGP GSSDKQEG<br>EWPTGLRLSRIGGIH* |
| 94 | MEYQHW **L**REAI SQLQASESPRRDAE <u>IL</u>LEHVTGKGRTFILAFGETQLTDEQC<br>QQLDALLTRRRDGEPIAHLTG VREF FSTPVWISQAQGI RAGP GSSDKQEGEW<br>PTGLRLSRIGGIH* |
| 91 | MEYQHWLREAI SQLQASESPRRDAE <u>IL</u>LEHVTGKGRTFILAFGETQLTDEQC<br>QQLDALLTRRRDGEPIAHLTG V FSTPVWISQAQGI RAGP GSSDKQEGEWPTG<br>LRLSRIGGIH* |
| 89 | MEYQHWLREAI SQLQASESPRRDAE <u>IL</u>LEHVTGKGRTFILAFGETQLTDEQC<br>QQLDALLTRRRDGEPIAHLTFSTPVWISQAQGI RAGP GSSDKQEGEWPTGLR<br>LSRIGGIH* |
| 84 | MEYQHWLREAI SQLQASESPRRDAE <u>IL</u>LEHVTGKGRTFILAFGETQLTDEQC<br>QQLDALLTRRRDGE P FSTPVWISQAQGI RAGP GSSDKQEGEWPTGLRLSRIG<br>GIH* |
| 80 | MEYQHWLREAI<u>SQLQASESPRRDAEILLEHVTGKGRTFILAFGETQLTDEQC</u><br>QQLDALLTRRRFSTPVWISQAQGIRAGPGSSDKQEGEWPTGLRLSRIGGIH* |
| 75 | MEYQHWLREAI<u>SQLQASESPRRDAEILLEHVTGKGRTFILAFGETQLTDEQC</u><br>QQLDALFSTPVWISQAQGIRAGPGSSDKQEGEWPTGLRLSRIGGIH* |
| 70 | MEYQHWLREAI<u>SQLQASESPRRDAEILLEHVTGKGRTFILAFGETQLTDEQC</u><br>QFSTPVWISQAQGIRAGPGSSDKQEGEWPTGLRLSRIGGIH* |
| 68 | MEYQHWLREAI<u>SQLQASESPRRDAEILLEHVTGKGRTFILAFGETQLTDEQF</u><br>STPVWISQAQGIRAGPGSSDKQEGEWPTGLRLSRIGGIH* |
| 66 | MEYQHWLREAI<u>SQLQASESPRRDAEILLEHVTGKGRTFILAFGETQLTDFST</u><br>PVWISQAQGIRAGPGSSDKQEGEWPTGLRLSRIGGIH* |
| 64 | MEYQHWLREAI<u>SQLQASESPRRDAEILLEHVTGKGRTFILAFGETQLFSTPV</u><br>WISQAQGIRAGPGSSDKQEGEWPTGLRLSRIGGIH* |
| 61 | MEYQHWLREAI<u>SQLQASESPRRDAEILLEHVTGKGRTFILAFGEFSTPVWIS</u><br>QAQGIRAGPGSSDKQEGEWPTGLRLSRIGGIH* |
| 59 | MEYQHWLREAI<u>SQLQASESPRRDAEILLEHVTGKGRTFILAFFSTPVWISQA</u><br>QGIRAGPGSSDKQEGEWPTGLRLSRIGGIH* |
| 56 | MEYQHWLREAI<u>SQLQASESPRRDAEILLEHVTGKGRTFIFSTPVWISQAQGI</u><br>RAGPGSSDKQEGEWPTGLRLSRIGGIH* |
| 52 | MEYQHWLREAI<u>SQLQASESPRRDAEILLEHVTGKGFSTPVWISQAQGIRAGP</u><br>GSSDKQEGEWPTGLRLSRIGGIH* |

List of FLN5-FLN6 constructs [1]. All constructs were modified to include a LepB derived C-terminal tail subsequent to the critical proline residue in the SecM(*Ec*) AP. The ‘length’ L^*^ corresponds to the number of residues between the C terminus of FLN5 and the critical proline residue at the C-terminal end of the SecM(Ec) AP. Non-folding controls with the Y719E mutation [1] were arrested with the SecM(*Ec*) AP, and were included for certain critical lengths used in the NMR study. Constructs 16, 11, and 1 were generated with only the stronger SecM(*Ec*, S→K) AP.

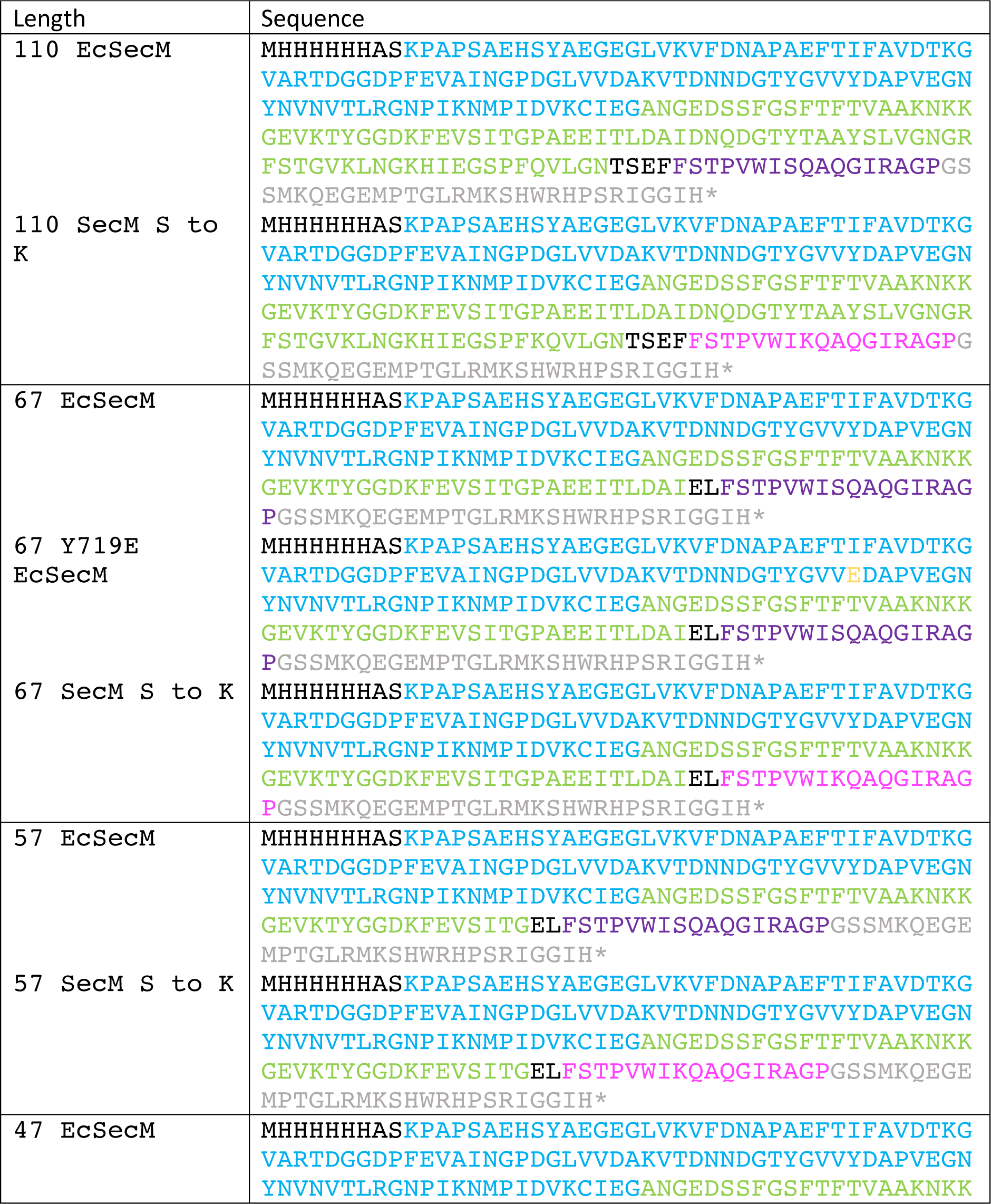
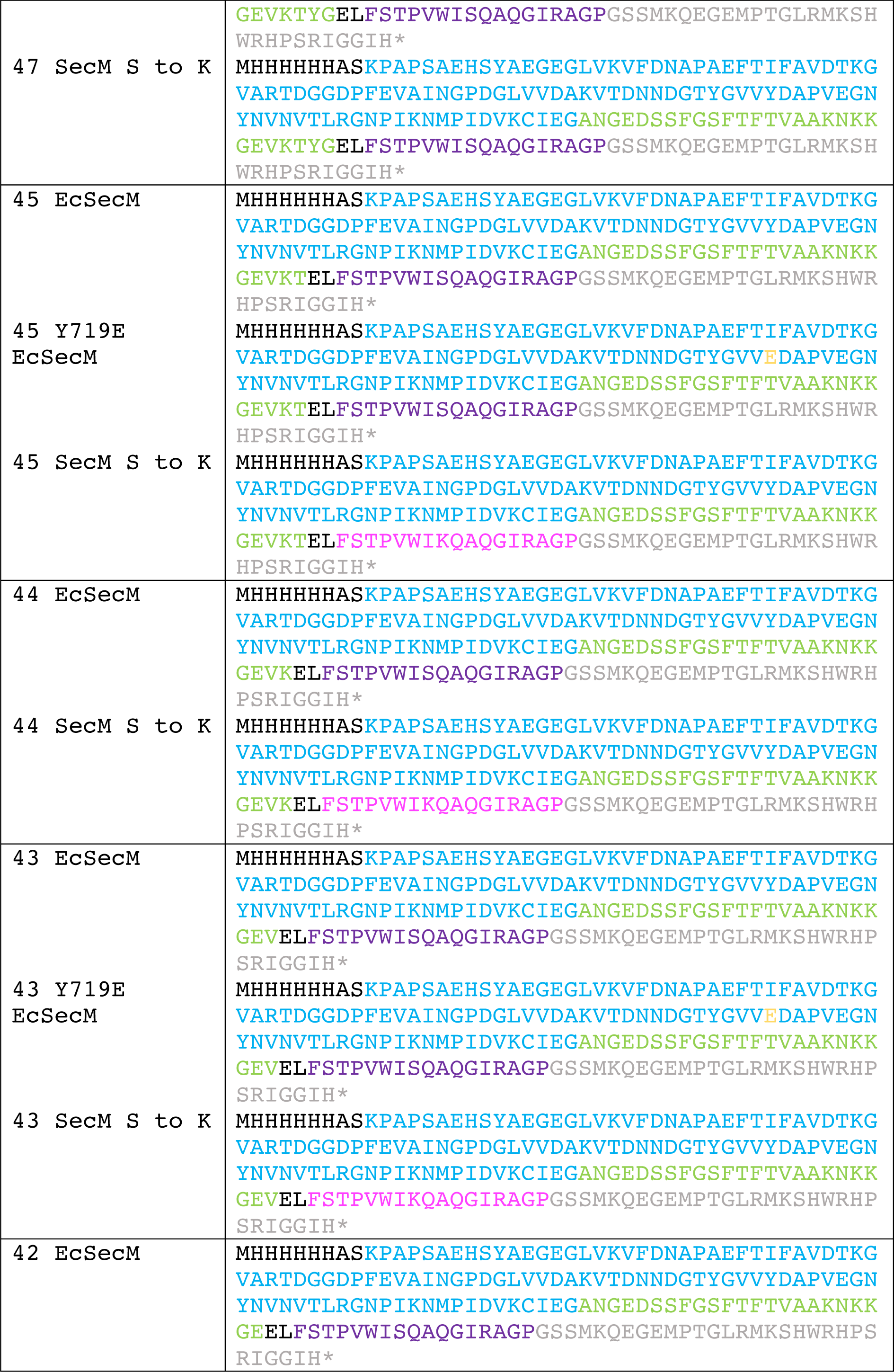
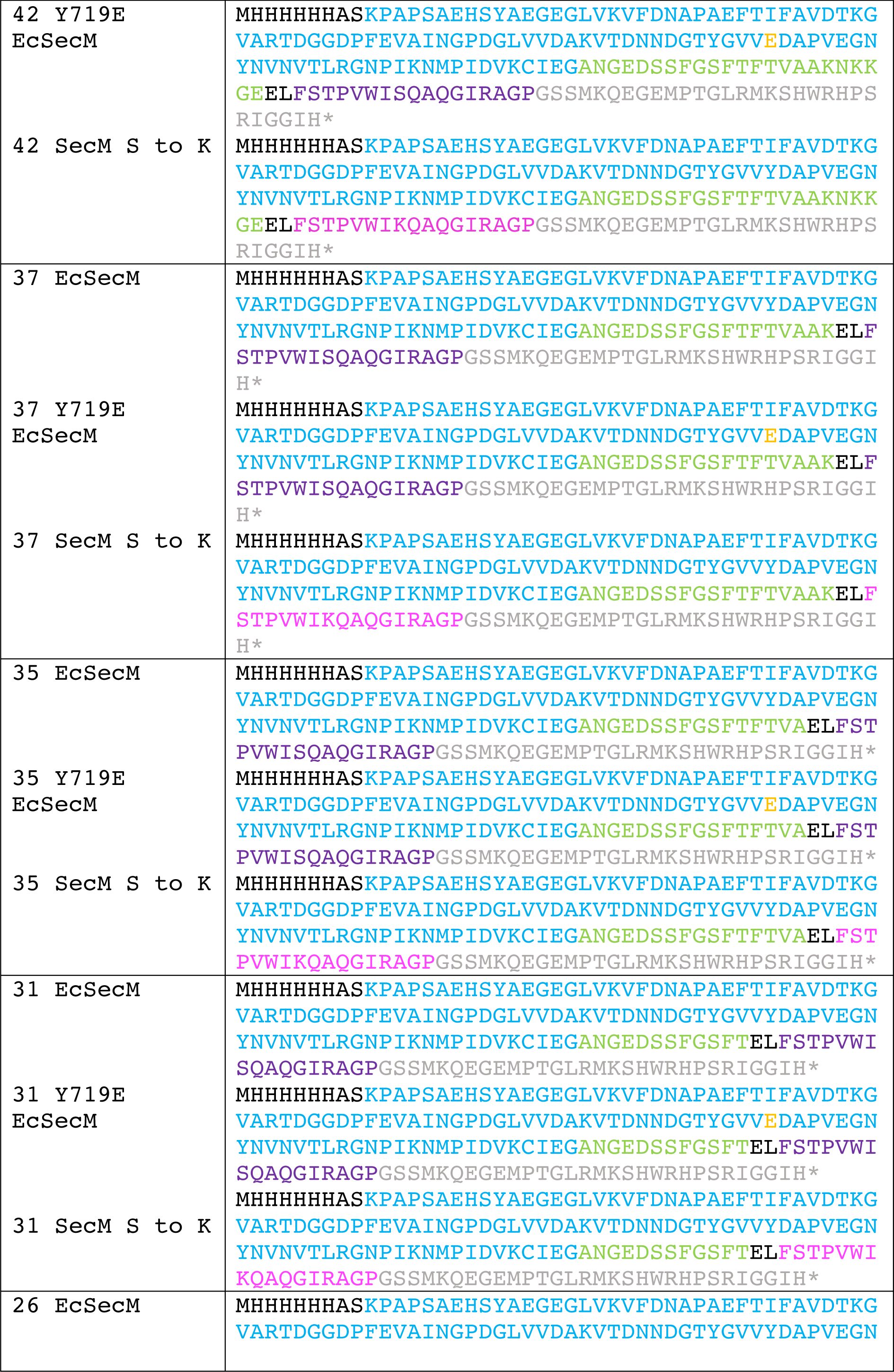
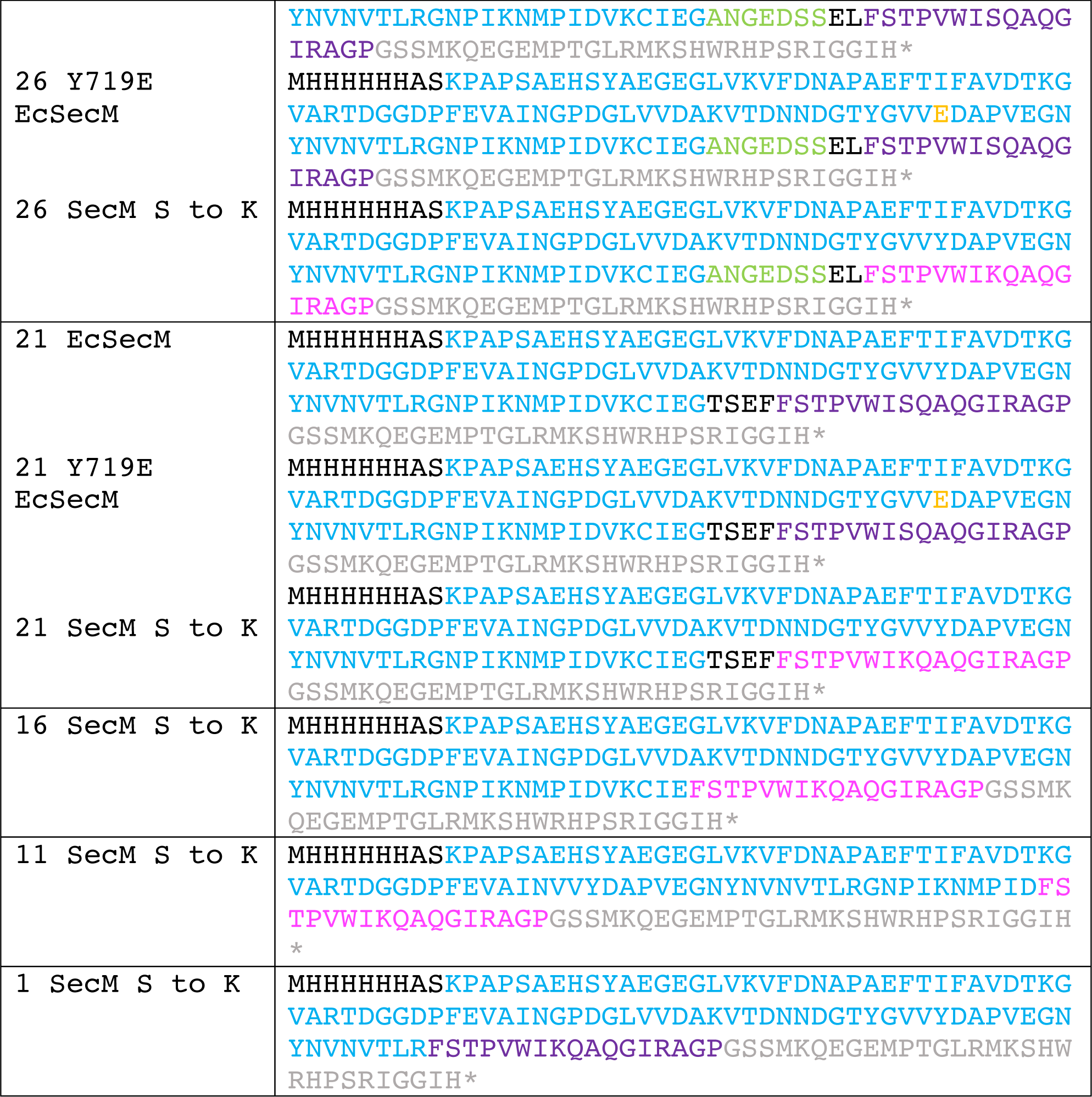

**Supplementary Fig. S1.**
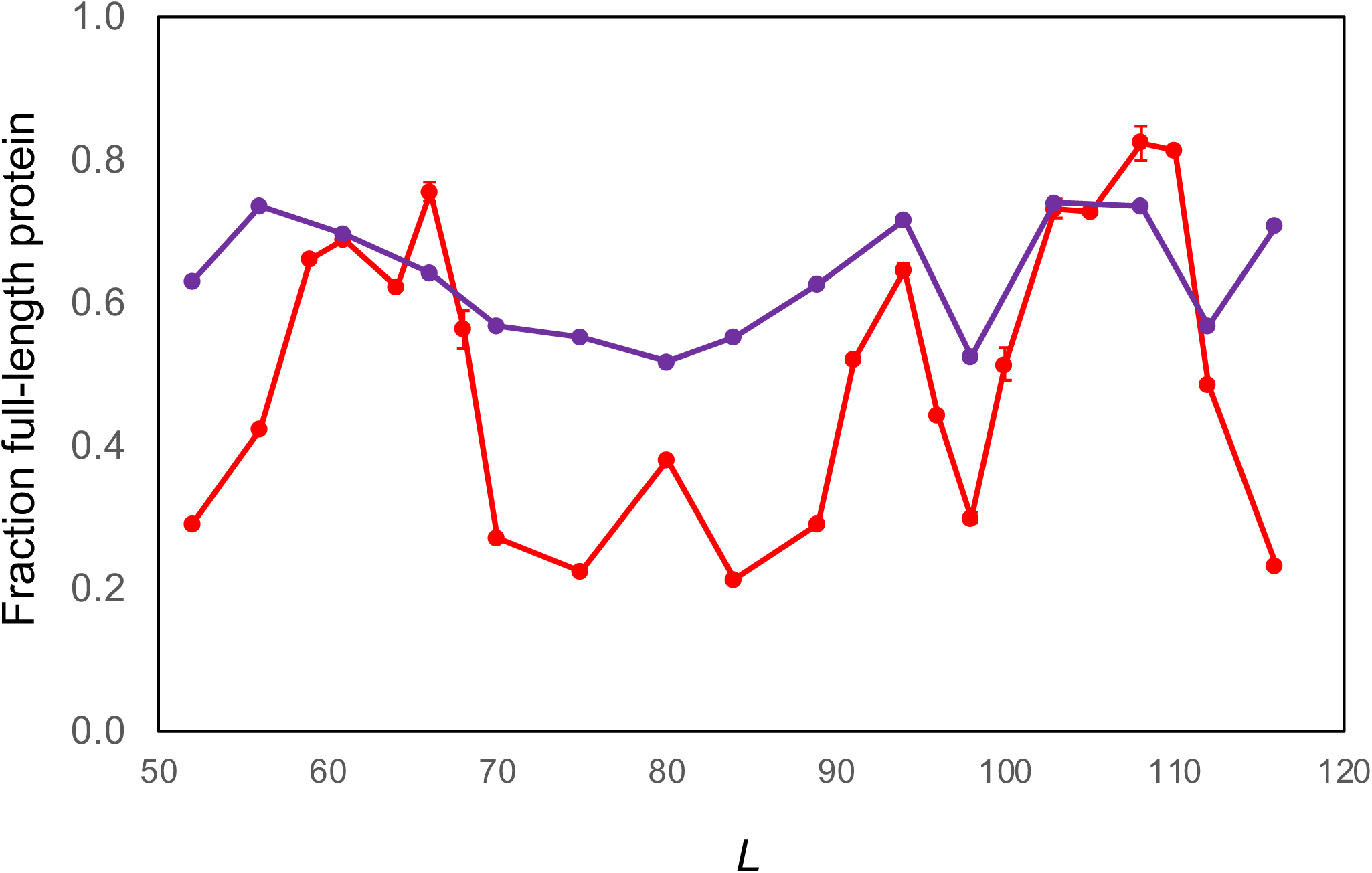
FPA profile obtained by *in vitro* translation in the *E. coli*-derived PURE *in vitro* transcription-translation system of wild-type HemK-NTD with the SecM(*Ec*) AP (red curve, c.f. Fig. 2b) and the SecM(*Ec-*HA) AP (purple curve) as the force sensor. The AP sequences can be found in Supplementary Table 1. All points represent the average of two measurements (all pairwise differences in *f*_*FL*_ ≤ 0.06), except where error bars are shown (SEM) in which cases 3 or 4 measurements were made.

**Supplementary Fig. S2.**
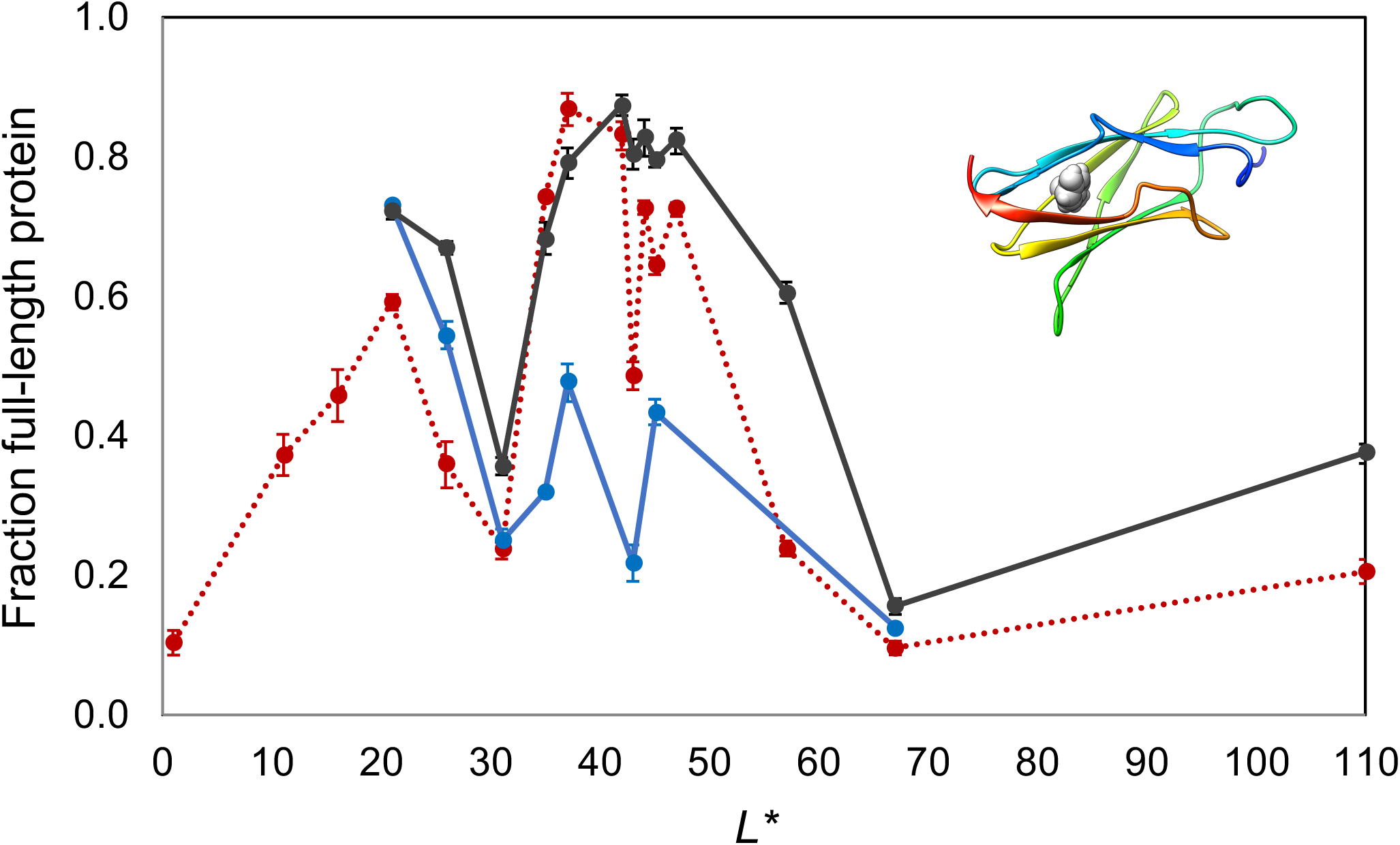
FPA of wild-type FLN5 (black curve) and of the non-folding mutant control Y^719^E (blue curve) with the SecM(*Ec*) AP as the force sensor, obtained by *in vitro* translation in the *E. coli*-derived PURE *in vitro* transcription-translation system. The force profile for FLN5 using the SecM(*Ec*, S→K) AP (dashed red curve, c.f. Fig. 3) is included for comparison. Error bars indicate SEM values. Inset is a ribbon representation of the backbone structure of FLN5 (PDB 2N62) with Y^719^ in spacefill.

**Supplementary Fig. S3.**
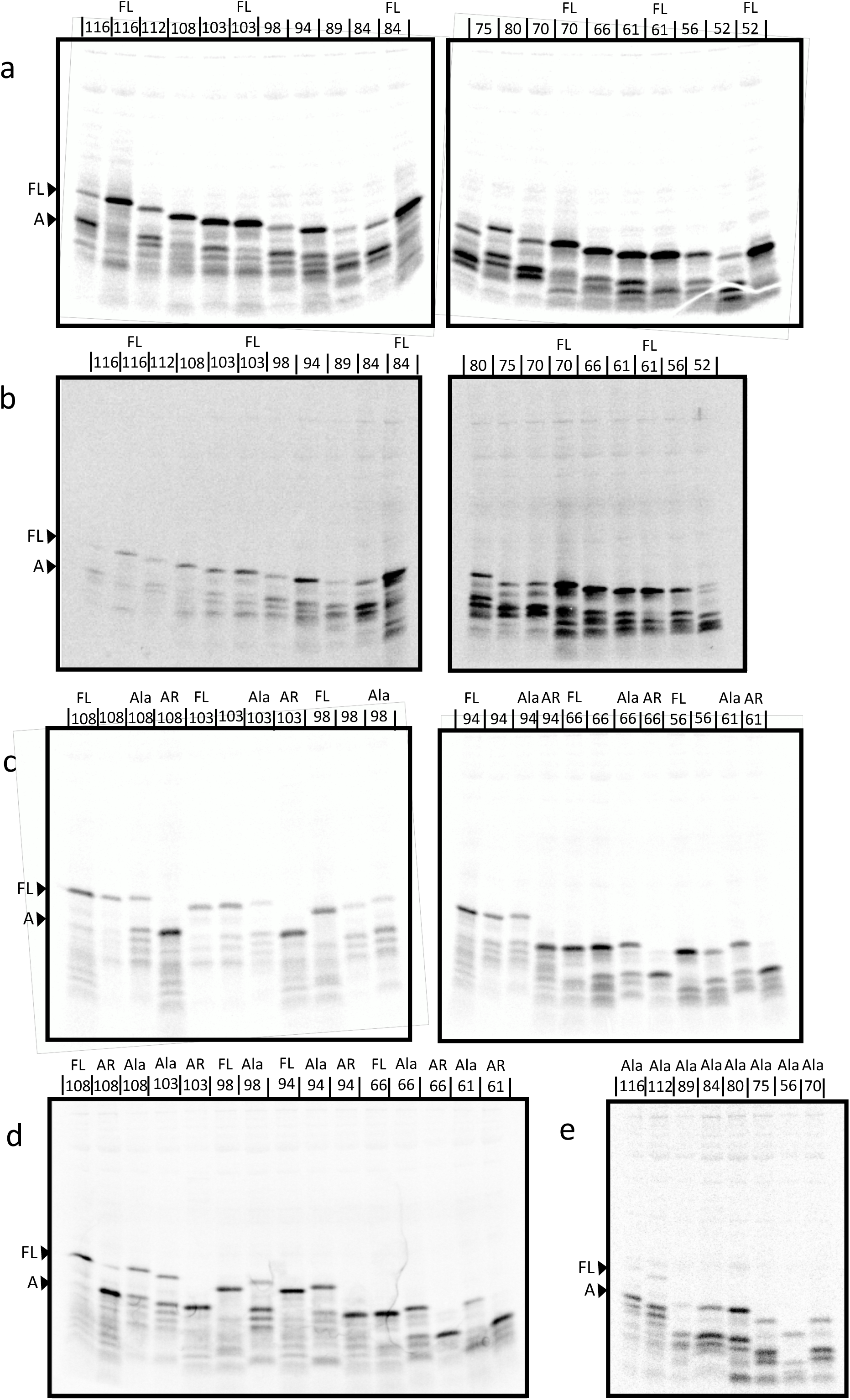
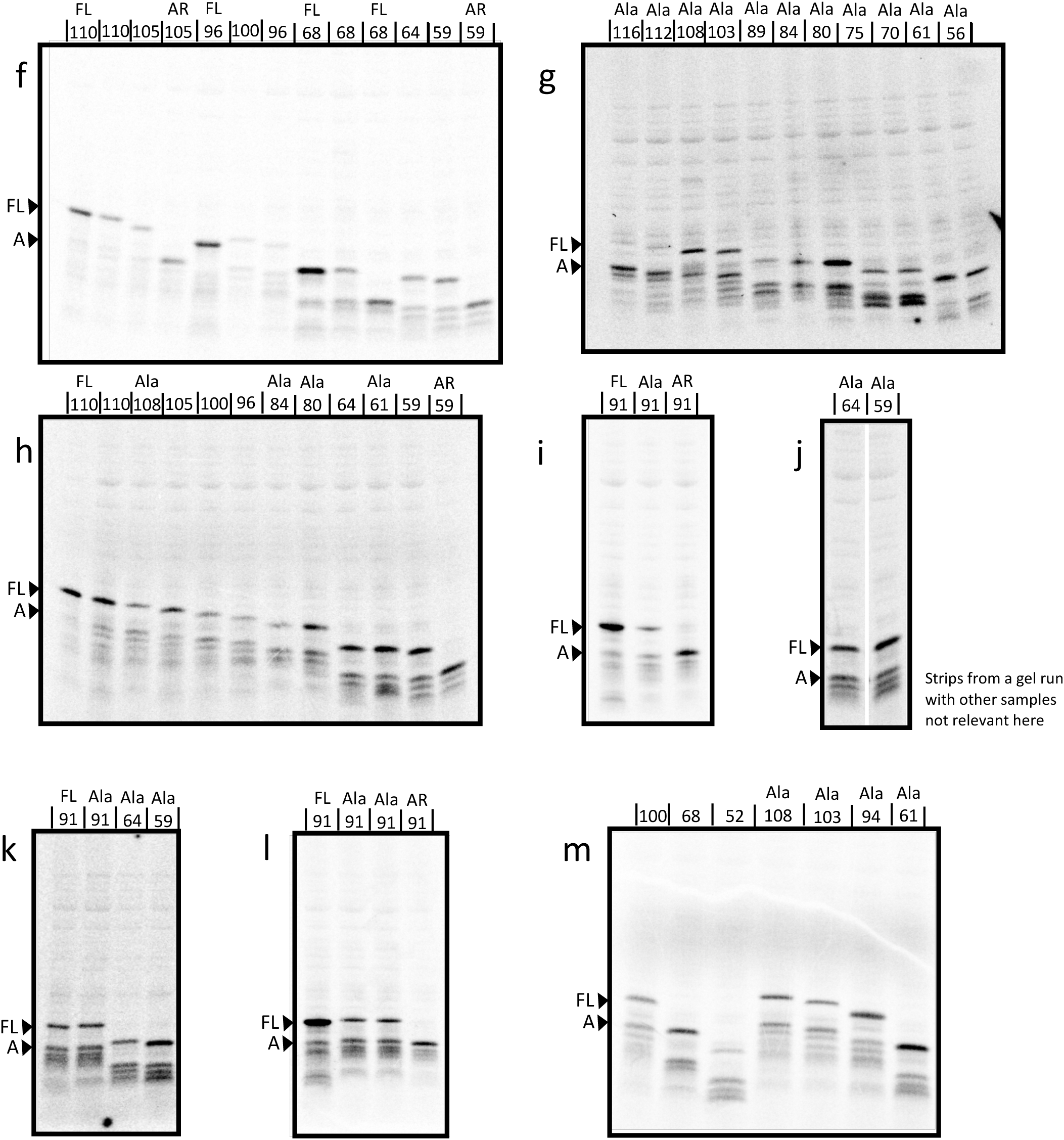
Replicates of autoradiographs of *in vitro* translated HemK-NTD constructs with the *E.coli* SecM(*Ec*) AP. *L* values are shown above each lane. SecM full-length and arrested control constructs are indicated by FL and AR respectively, and Leu^27,28^Ala mutants are indicated by Ala.

**Supplementary Fig. S4.**
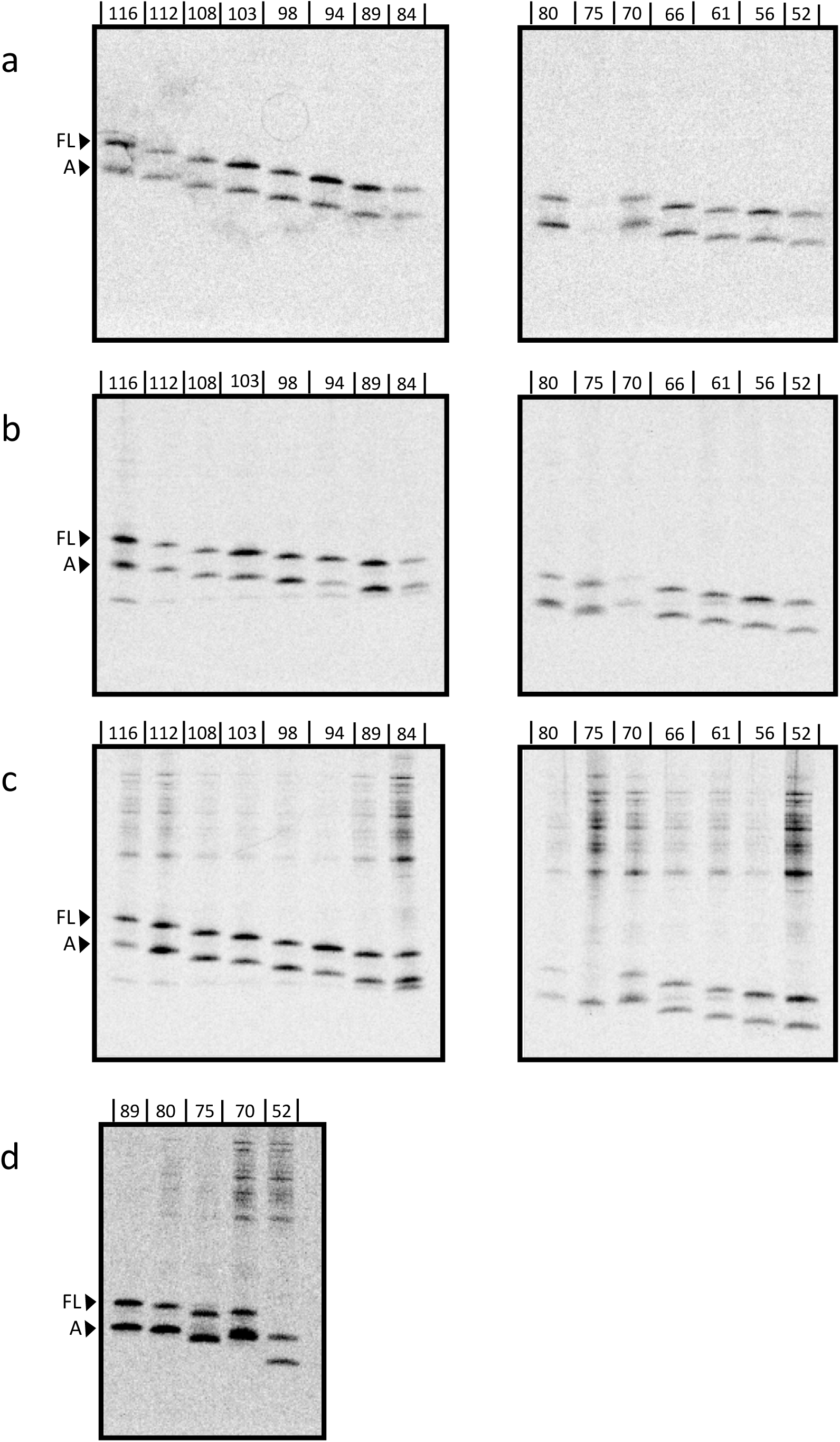
Triplicates of autoradiographs of *in vivo* translated HemK-NTD constructs stalled with the *E.coli* SecM(*Ec*-HA) AP. *L* values are shown above each lane.

**Supplementary Fig. S5.**
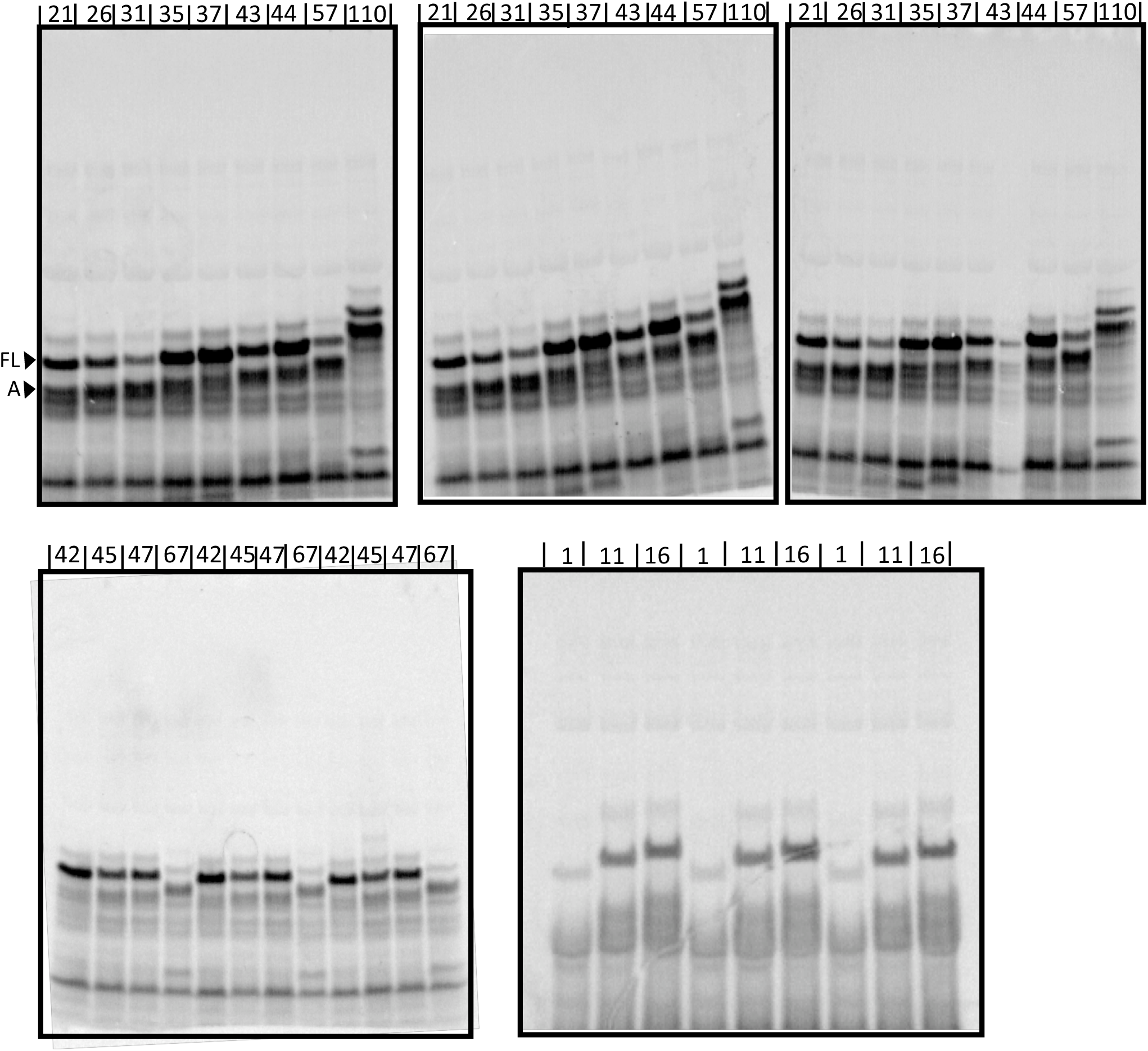
Triplicates of autoradiographs of FLN5-FLN6 constructs with the SecM(*Ec*, S→K) AP. *L\** values are shown above each lane.

**Supplementary Fig. S6.**
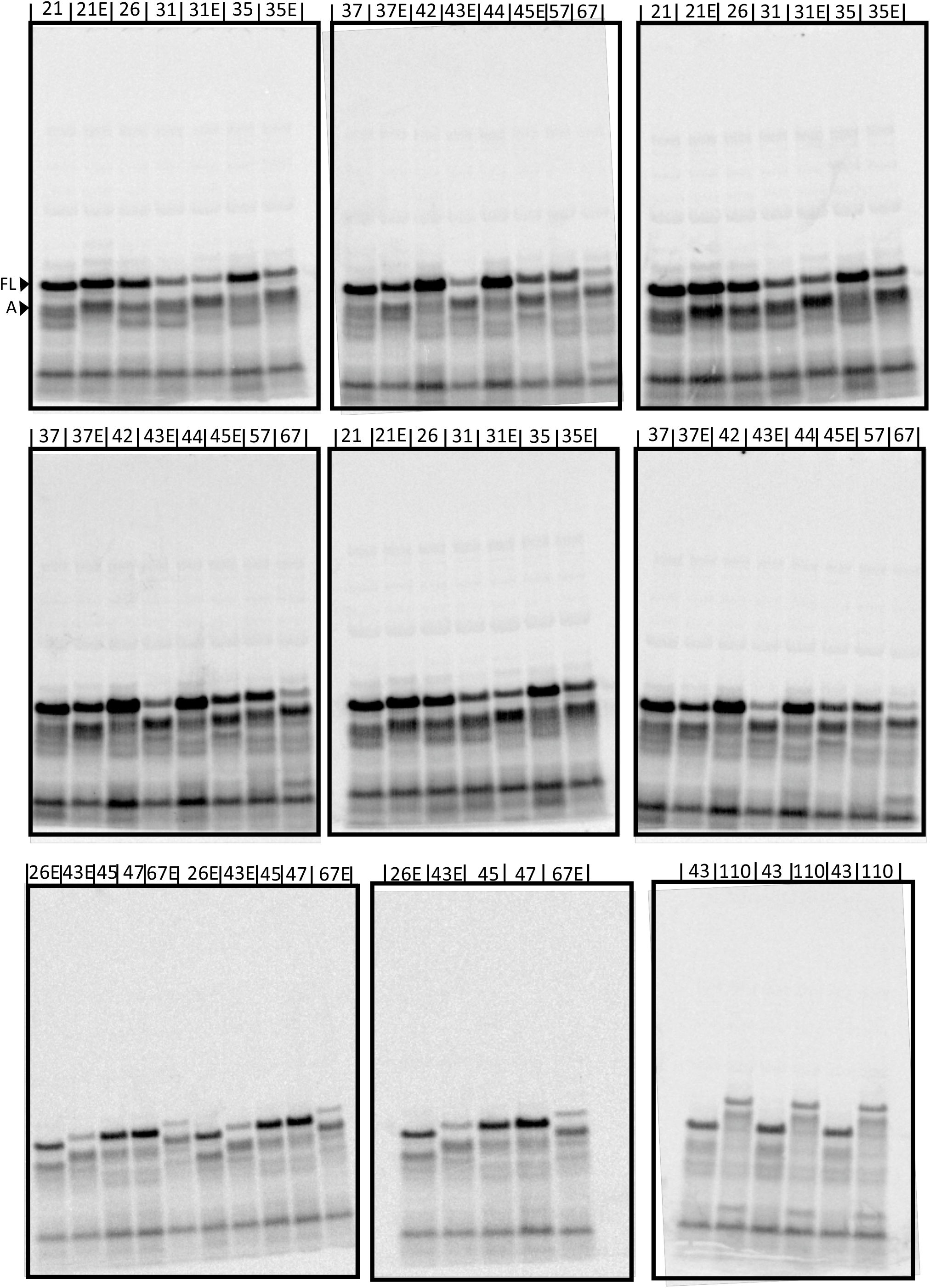
Triplicates of autoradiographs of FLN5-FLN6 constructs with the SecM(*Ec*) AP. Non-folding controls (Y^719^E mutation) indicated by”E” are included for certain lengths. *L\** values are shown above each lane.

